# Variable performances of commercial eDNA inventories challenge their use for surveying stream fish communities

**DOI:** 10.64898/2026.03.15.711554

**Authors:** JM Roussel, E. Quéméré, B. Bonnet, R. Covain, O. Dézerald, G. Lassalle, PY. Le Bail, E.J. Petit, G. Pottier, G. Quartarollo, R. Vigouroux, H. Lalagüe

## Abstract

1. Environmental DNA (eDNA) metabarcoding of water samples is increasingly used to detect fish species in streams. Several studies have concluded that it can outperform traditional inventory methods and recommend using it at large scales for fish-based ecological assessments. However, there is no standard protocol that can guarantee sufficient detection rates and repeatability, despite companies offering an extensive range of analyses.

2. We compared eDNA metabarcoding performed by four companies. Following their guidelines, samples were collected in a small tropical stream in the Maroni River (French Guiana) that hosts a species-rich fish community. We compared their inventories to each other and to a list of species captured during an extensive fish inventory performed immediately after sampling eDNA, as well as to current data on the species’ distributions.

3. The number of species detected by eDNA metabarcoding ranged from 5 to 48 among the companies, but these inventories contained many inaccuracies. All companies combined, 63 species were detected, of which 10 (16%) had never been reported in the Maroni River. The extensive inventory identified 50 species in the local fish community, of which 16-46 were not detected by eDNA metabarcoding (i.e. false negative detection rate of 32%-92% among the companies).

4. Reanalysis of raw sequencing data decreased differences among companies greatly, highlighting the importance of using a comprehensive and accurate DNA barcode database to assign species. Dissimilarity indices, calculated to compare the local fish community (based on presence/absence or fish catches) to eDNA detection, revealed large differences regardless of the company.

5. Summary and applications. The large percentage of species not detected by eDNA metabarcoding of water samples could strongly bias fish-diversity inventories in streams that host species-rich communities. This issue is not well documented in the literature, and we recommend that similar studies in the future focus on other stream contexts. The large differences between commercial eDNA inventories and the local fish community challenge the use of eDNA metabarcoding for fish-based ecological assessments of streams. The variable performance of eDNA companies indicates the need for a standard protocol and access to a comprehensive DNA database before beginning large-scale eDNA programmes.

**Highlights:**

- eDNA metabarcoding of water samples is widely used to detect species in streams
- Detection performances of 4 private companies were compared to an exhaustive fish inventory
- The number of undetected species varies from 32 to 92% depending on the company
- Such discrepancies challenge the use of eDNA for fish-based ecological assessments

## Introduction

Environmental DNA (eDNA) refers to the genetic material that organisms leave behind in the environment. It is used to detect the presence of species in environmental samples, which allows biodiversity to be studied with an unprecedented potential for action. After the pioneering study two decades ago (Ficetola et al., 2008), eDNA was soon promoted as the ultimate biomonitoring tool (Baird & Hajibabaei, 2012). Inexpensive, accurate, simple to use in the field, and ethically-based, eDNA metabarcoding was quickly set to enhance, or even replace, traditional methods based on capturing or observing organisms, thus helping policy making and future practices better prevent further biodiversity loss (Taberlet et al., 2012). This has resulted in a large amount of research and development, with nearly 1000 articles about the use of eDNA for biomonitoring published over the past decade (Blackman et al., 2024). When no previous knowledge about a community is available, eDNA metabarcoding has become the most widely used approach, as it can identify and inventory species using molecular primers that target broad taxonomic groups (Cordier et al., 2020; Deiner et al., 2017). Many studies have compared eDNA metabarcoding to traditional methods, and they often reported that the former performed as well as or better overall than the latter (*e.g.* Andres et al., 2023; Bonk et al., 2024; Czeglédi et al., 2021; Jannel et al., 2024; Keck et al., 2022; Liu et al., 2024). This has led many scientists and government agencies to recommend large-scale use of eDNA metabarcoding in monitoring programmes, especially for freshwater ecosystems (*e.g.* Keck et al., 2022; Kelly et al., 2024; Pawlowski et al., 2018; Stein et al., 2023). However, closer examination of studies that compared it to traditional methods for freshwater ecosystems revealed that its performance may be exaggerated due to several unresolved problems, and that key challenges remain to be addressed before it is routinely used for biomonitoring.

First, eDNA metabarcoding of water samples is usually compared to traditional sampling protocols, which also inventory only some of the species present. Few field studies have combined eDNA metabarcoding with exhaustive inventories of communities to explore the former’s limitations (but see Blabolil et al., 2021; Di Muri et al., 2020), and the former’s seemingly better performance should not mask its potential lack of completeness in the field. In general, the absence of a species is difficult to confirm unless extensive sampling is performed, and eDNA metabarcoding is no exception. Some organisms release DNA fragments into aquatic environments, depending on their taxonomy, metabolism, and ontogenetic stage (Múrria et al., 2024; Poyntz-Wright et al., 2024; Tréguier et al., 2014; Verdier et al., 2024; Yates et al., 2019; Yates et al., 2021a; Yates et al., 2021b). Thus, a species may be present, but its eDNA concentrations lie below the detection threshold. The eDNA metabarcoding protocol includes several key steps – water sampling in the field, DNA extraction and amplification in the laboratory, DNA sequencing, and bioinformatic analysis (Deiner et al., 2017) – during which the eDNA signal of a species may be lost. The risk of misinterpreting non-detection of eDNA as true species absence (*i.e.* false negative detection) exists and must be carefully considered when using eDNA (Lacoursière-Roussel & Deiner, 2021).

Second, several studies have found large differences between species lists identified using eDNA metabarcoding and traditional methods, especially at the local scale (Brantschen & Altermatt, 2024; Cote et al., 2023; Keck et al., 2022; Kelly, 2019; Liu et al., 2024). Besides animal movement, water currents can transport DNA fragments over long distances (Civade et al., 2016; Deiner & Altermatt, 2014; Harrison et al., 2019). Notably, rivers act as “conveyor belts” of biodiversity information (Deiner et al., 2016; Littlefair et al., 2023) in which the area represented by an eDNA sample depends on the persistence, shedding, decay, and downstream transport of DNA, all of which vary greatly and are influenced by complex environmental interactions among discharge, exposure to ultraviolet light, substrate composition, biofilm and microbial activity, and suspended-sediment characteristics (Condachou et al., 2025; Deiner & Altermatt, 2014; Harrison et al., 2019; Hervé et al., 2023; Mauvisseau et al., 2022; McKnight et al., 2024; Nevers et al., 2020; Scriver et al., 2023; Shogren et al., 2017; Snyder et al., 2023; Turner et al., 2015). Animal waste can also spread eDNA in the environment (Inoue et al., 2023; Stoeckle et al., 2016), such as when a predator excretes the remains of its prey outside of the prey’s foraging habitats (Guilfoyle & Schultz, 2017; Merkes et al., 2014). These factors imply that detecting a species’ DNA in a geolocated environmental sample does not necessarily indicate its actual presence at that location, and that eDNA samples may represent a different spatial scale than those of traditional methods. Unlike methods based on capturing animals, eDNA introduces a new source of error – false positive detection – whose extent remains largely unexplored.

Third, the influence of false negatives and positives on sampling design and statistical procedures is not sufficiently recognised. The development of eDNA metabarcoding as a tool in biodiversity studies has been driven by technological revolutions, especially the advent of high-throughput sequencing technologies and the initial realisation that DNA can be extracted from environmental samples (Ficetola et al., 2008). Despite the early recommendation of Yoccoz (2012) that the eDNA community should collaborate with quantitative ecologists, relevant sampling designs are rarely adopted, which may hinder their ability to provide information about the state of and changes in biodiversity (Yoccoz et al., 2001). A good indicator for eDNA inventories is the ratio of biological replicates (i.e. water samples) to technical replicates (i.e. Polymerase Chain Reaction; PCR), which is usually low. Most eDNA protocols prefer PCR replicates to water-sample replicates, but both are needed to assess potential sources of errors correctly (Guillera-Arroita et al., 2017; Lahoz-Montfort et al., 2016). This disregard for statistical issues is especially prevalent for aquatic environments, as demonstrated by recent reviews that did not address them (*e.g.* Bruce et al., 2021; Bunholie et al., 2023; Xing et al., 2022). Most eDNA developments have focused on increasing sample volumes rather than the number of biological replicates (Blackman et al., 2024), and using statistical models to quantify errors remains the exception (15 of the 431 studies reviewed by Mathieu et al. (2020)).

The increasing demand for eDNA-based tools to monitor biodiversity has spurred the worldwide emergence in recent years of many companies that provide eDNA metabarcoding, including “turnkey” eDNA solutions for species inventories. They provide clients (i.e. government agencies, NGOs, and research institutions) with user-friendly sampling kits that allow the clients to filter water samples themselves in the field. The companies process the samples in the laboratory, perform the metabarcoding, and provide a list of the taxa detected, usually with little information about the reliability of their results. Studies have highlighted the lack of minimum justifications for eDNA methods and sampling design (Blackman et al., 2024; Mathieu et al., 2020), especially the volume filtered, filter pore size, and numbers of replicates, which vary greatly among water eDNA studies (Bunholi et al., 2023; Shu et al., 2020; Xing et al., 2022). Moreover, despite the development of large curated barcode databases (e.g. BOLD, SILVA, MitoFish) over the past few decades, their lack of comprehensiveness and accuracy remains a major challenge for implementing biomonitoring programmes correctly (Weigand et al., 2019), especially for species-rich communities (Keck et al., 2023; Marques et al., 2021). The development of private databases poses accessibility problems for users, increases the risk of incorrect taxonomic assignment, and could result in many false positive and/or false negative detections in eDNA inventories (Jackman et al., 2021). Companies vary greatly in pricing, methodological transparency, and data-sharing policies, which makes it difficult for non-academic users to select among them. It also poses challenges for comparing eDNA inventories among companies, especially for long-term biomonitoring programmes.

Following promising scientific developments, eDNA inventories are increasingly expected to be used to monitor biodiversity, including for regulatory surveillance of aquatic environments (Hering et al., 2018; Pawlowski et al., 2018; Pont et al., 2021). The number of tendering processes is likely to increase in the future, but the accuracy with which commercial eDNA inventories are able to describe fish communities at a given location is unknown. In the present study, we compared four companies that perform eDNA metabarcoding of water samples to assess the taxonomic composition of fish communities. The study was performed using a species-rich stream in French Guiana to challenge the performance of eDNA metabarcoding. After carefully following each company’s guidelines for sampling water and preserving the samples, we sent the samples to each company without disclosing the study’s objective and the exact sampling location. The species lists that each company provided were compared to (i) an exhaustive inventory of fish species performed at the sampling site immediately after sampling the water and (ii) current data on fish distribution in French Guiana. False negative and false positive eDNA detection rates and fish-community composition indices were calculated and compared among the companies. Finally, raw sequencing data supplied by three of the companies were reanalysed using a standard bioinformatic pipeline and a recently published barcode database of fish species in French Guiana. We then assessed how the data analysis may have influenced differences among the companies.

## Methods

### Study site and eDNA sampling

The Bastien stream flows into the Maroni River in French Guiana (Supplementary Material S1). eDNA was sampled on 18 March 2024 in a 2.5 km tributary of the Bastien stream (5.27048° N, 54.234217° W; Supplementary Materials S1 and S2). Water temperature (26.1°C), conductivity (32.1 µS.cm^-1^), turbidity (2.83 NTU), pH (6.02), and dissolved oxygen (6.52 mg.L^-1^, 79.8% saturation) were recorded using a multi-parameter probe (Tripod Ponsel Aqualabo). Stream discharge (15 L.s^-1^) was estimated from water depth and velocity measured across the channel section. The four eDNA companies selected are internationally known. Following their methodological guidelines, stream water was pumped in running water to ensure that it was sufficiently mixed, and then filtered in the field using sampling kits provided by each company. Differences in the number of filters used, pore size, filtration times, and volumes filtered differed (Supplementary Material S3) revealed diverse sampling strategies among companies. After sampling, the filters were stored according to each company’s guidelines and placed in a cool box for transport to the laboratory.

eDNA samples were sent to the companies for eDNA metabarcoding, without revealing the objective of the study or the sampling locations, to avoid influencing the processing of samples. We asked them only to inventory the fish species detected in the samples. As the companies provided few details about their laboratory workflows, bioinformatic pipelines, or statistical treatments, we were not able to compare these aspects directly. Three companies stated that they used one primer pair, either 12S-Teleo (64 bp) (Valentini et al., 2016) (*Teleo*) or 12S-MiFish-U (163-185 bp) (Miya et al., 2015) (*MiFish*), while the fourth stated that it used several primer pairs (targeting 12S, COI and 16S regions), but did not specify which ones. Three companies analysed two biological replicates, while the fourth analysed three (Supplementary Material S3). The number of technical (PCR) replicates per sample varied greatly: from 1-12 for the three companies that provided this information (Supplementary Material S3). Finally, three of the four companies agreed to send us their raw sequencing data (Illumina FastQ files). The companies have been anonymised to avoid influencing their commercial activities.

### Exhaustive inventory of the local fish community

Immediately after filtering the water, a barrier net (4 mm mesh) was placed across the channel immediately on the water-pumping point (Supplementary Material S2). The bottom of the net was buried in the substrate to prevent fish from escaping underneath. Additional barrier nets were installed upstream to enclose a 100 m long section and prevent fish from escaping during the inventory. The mean channel width in the section was 225 cm (range: 150-375 cm). The habitat was a mean of 16.9 cm deep (range: 2-50 cm). The stream bottom consisted mainly of pebbles, sand, and silt, with a few macrophytes (*Thurnia sphaerocephala*). We inventoried the fish community in the enclosed section exhaustively using electrofishing. The electrofishing device (described by Pottier et al. (2019)) was set to deliver 1000 V (DC). The anode was a 350 mm diameter aluminium ring, and the cathode was a double copper braid (3000 mm long × 10 mm wide × 1 mm thick). Given the ambient conductivity, these settings ensured effective fish capture while maintaining animal welfare (Pottier et al. (2019) for details). The fisherman who managed the anode was assisted by two people equipped with dip nets to capture the fish in the anode’s electric field. The entire water surface in the enclosed section was explored from downstream to upstream, four times in a row, to maximise fish capture.

After capture, each fish was identified to species using the latest taxonomic knowledge about fish in French Guiana (Le Bail et al., 2012; Le Bail et al., 2026). Fish were measured (standard length) to the nearest mm and then gradually released downstream of the sampling section. We applied allometric length-weight relations based on a historical database of fish captures in French Guiana (https://eseweb.rennes.inrae.fr/RezoFleuve/) (Allard et al., 2015) to estimate the weight of each individual, and summed the weights to calculate the biomass of each fish species in the community. Hereafter, “local fish community” refers to the abundances and biomasses of species captured in the 100 m enclosed section.

### False positive and false negative eDNA detection rates

The names of taxa that the companies provided were standardised according to Le Bail et al. (2026) and Fricke et al. (2025). The number of taxa detected at the class, order, genus, and species levels were counted. Companies were not penalised for incorrect spelling or if the taxonomy at the species level was still debated by taxonomists. However, taxa reported only to genus, family, order, or class were excluded when taxonomic knowledge could have been used to identify them to species. Each company’s list was analysed independently before being combined into a single list of all species detected by eDNA (hereafter, the “molecular community”). We created Venn diagrams to compare species lists among companies using the *eulerr* package (Larsson, 2024) of R software (R Core Team, 2025). We also used current knowledge about fish distribution to compare each company’s species list to those in the Maroni River catchment, other large rivers in French Guiana, and elsewhere in South America (Le Bail et al., 2026).

As no company provided information about taxon detection across biological and technical replicates, we used the fish captures as unambiguous detections (i.e. no false positive detections) (Guillera-Arroita et al., 2017; Lahoz-Montfort et al., 2016; Miller et al., 2011). Species in the fish community that were not detected by eDNA metabarcoding were classified as false negative detections, whose rates were calculated for each company by dividing the number of undetected species by the number of species captured in the local fish community. The same calculation was used for the molecular community. Conversely, species detected using eDNA metabarcoding that were absent from the local fish community were classified as false positive detections, whose rates were calculated for each company and for the molecular community by dividing the number of species detected using eDNA barcoding but not captured in the local fish community by the total number of species detected using eDNA barcoding.

### Reanalysis of companies’ raw sequencing data

To decrease potential biases related to unequal access to DNA databases among companies and to highlight differences caused by field sampling and/or laboratory processing, we reanalysed the raw sequencing data provided by three of the companies using a standard bioinformatic pipeline. Forward and reverse reads were assembled using the *illuminapairend* function of the *OBITools* v1.2.11 package (Boyer et al., 2016), and the resulting sequences were trimmed for primers using the cutadapt v1.8.3 program (Martin, 2011). Strictly identical sequences were clustered using the *obiuniq* function (dereplication step). Amplicon sequence variants represented by fewer than 10 reads were removed from the dataset. Additionally, sequence lengths were filtered for each primer set: 40-100 and 140-200 bp for *Teleo* and *MiFish*, respectively. Taxa were assigned using BLASTn (NCBI BLAST+ v2.10.0) compared to a curated reference database (Brosse et al., 2026; NCBI Accession Number: PX415477 - PX417033), with a minimum similarity threshold of 97% and 98%, an e-value cut-off of 1e^-70^ and 1e^-20^, and a minimum alignment coverage of 87% and 80% for *MiFish* and *Teleo*, respectively. When several equally good hits occurred, taxa were assigned to the lowest common ancestor. The resulting species lists were then used to recalculate false positive and false negative detection rates using the procedure described previously.

### Statistical analysis of composition of the local fish community

Species lists provided by the four companies were used to generate presence/absence data, while read counts from reanalysing companies’ raw sequencing data were used as a proxy for fish abundance (companies A, B, and D only). We compared eDNA (for each company individually or combined) to the local fish community using four dissimilarity indices: Jaccard and Raup-Crick for presence/absence, Bray-Curtis and Horn-Morisita for abundance data. Raup-Crick (Chase et al., 2011) and Horn-Morisita (Chao et al., 2006), which considers the co-occurrence of rare and unshared species among communities more than the Jaccard and the Bray-Curtis indices do, are more robust to differences in species composition and sample size. All dissimilarity indices among communities were calculated using the *vegan* package (Oksanen et al., 2024) and then visualised via principal coordinate analysis, with Cailliez-transformation when needed, using the *ade4* package (Dray & Dufour, 2007).

Finally, we tested whether the abundance and biomass of species in the local fish community influenced their detection by eDNA barcoding based on the reanalysis of raw sequencing data. Welch’s t-tests (unequal variance, two samples) were used to compare abundance and biomass of the species detected, or not, by eDNA metabarcoding. Linear regressions between abundance (or biomass) of species in the local fish community and read counts were calculated, and their significance was assessed using F-tests. All analyses were performed in R.

## Results

### Exhaustive inventory of the local fish community

The 446 fish captured via electrofishing belonged to 6 orders, 22 families, 40 genera, and 50 species (Supplementary Material S4). The local fish community was dominated by species with low abundance: 56% of the species were represented by five individuals or fewer, including nine species represented by only one individual (Supplementary Material S4). Conversely, five species were represented by more than 20 individuals (Supplementary Material S4 and S5). The total biomass captured was 2.037 kg for all species combined, of which 54% consisted of individuals weighing less than 10 g. The total biomass of 11 species exceeded 50 g each, being largest (428 g) for the predator *Hoplias malabaricus* (8 individuals).

### Assessment of species lists provided by the eDNA companies

The species list reported by each company (Table 1) contained several taxonomic inaccuracies, including 4 assignments to class (*Actinotperygii*), 1 to order, 5 to family, and 37 to genus. The companies reported varying levels of taxonomic resolution: the percentage of taxa identified to species was highest for company A (94%), followed by companies C (61%), B (59%), and D (24%). After removing inaccurate detections and standardising the taxonomy, the number of species detected by eDNA ranged from 5-48 among the four companies (Table 1). The molecular community contained 63 species. Sequencing depth varied greatly among companies, with company A producing 12 times as many reads as company B (Supplementary Table S3). Company A detected the most species and had the most read counts in its raw sequencing data. See Supplementary Material S6 for the complete species list reported by each company.

**Table 1.**
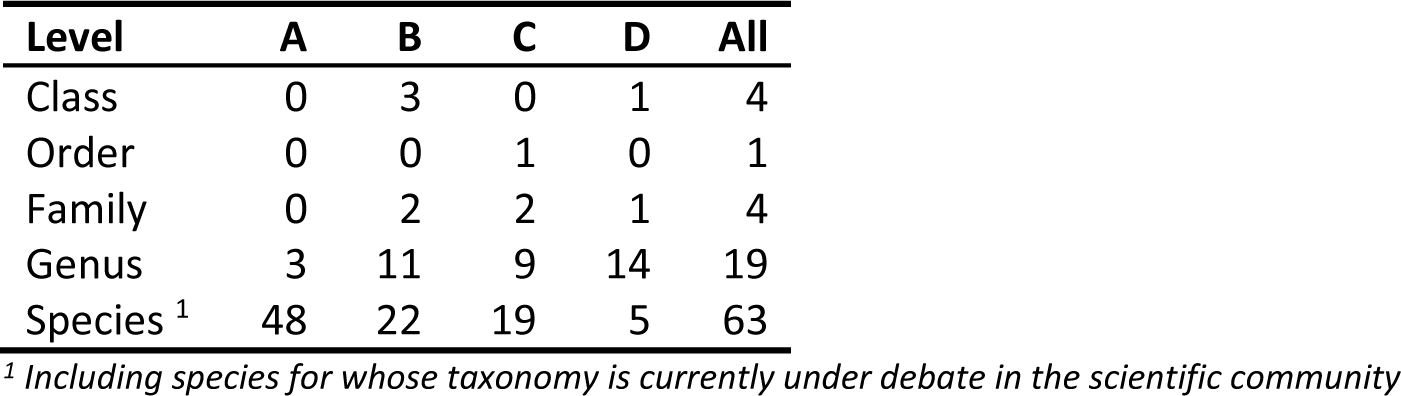
Number of taxa reported by eDNA companies (A, B, C, D and all combined) at the class, order, family, genus, and species levels.

The percentage of species detected by eDNA metabarcoding that were actually in the local fish community ranged from 41%-80% among the companies (Figure 1). The molecular community included 57% of the species actually in the local fish community, along with additional species known to inhabit the Maroni River catchment (27%), other large rivers in French Guiana (3%), or elsewhere in South America (13%). Company A detected 30 species that the other companies did not (Figure 1), most of which were species known to inhabit the Maroni River catchment (Supplementary Material S6). Company B detected 13 unique species, 7 of which had never been reported in the Maroni River catchment or other large rivers in French Guiana. Companies A and C shared the most detections in common: 17 species (Figure 1). Ultimately, only four species were detected by all four eDNA companies (Figure 1).

**Figure 1.**
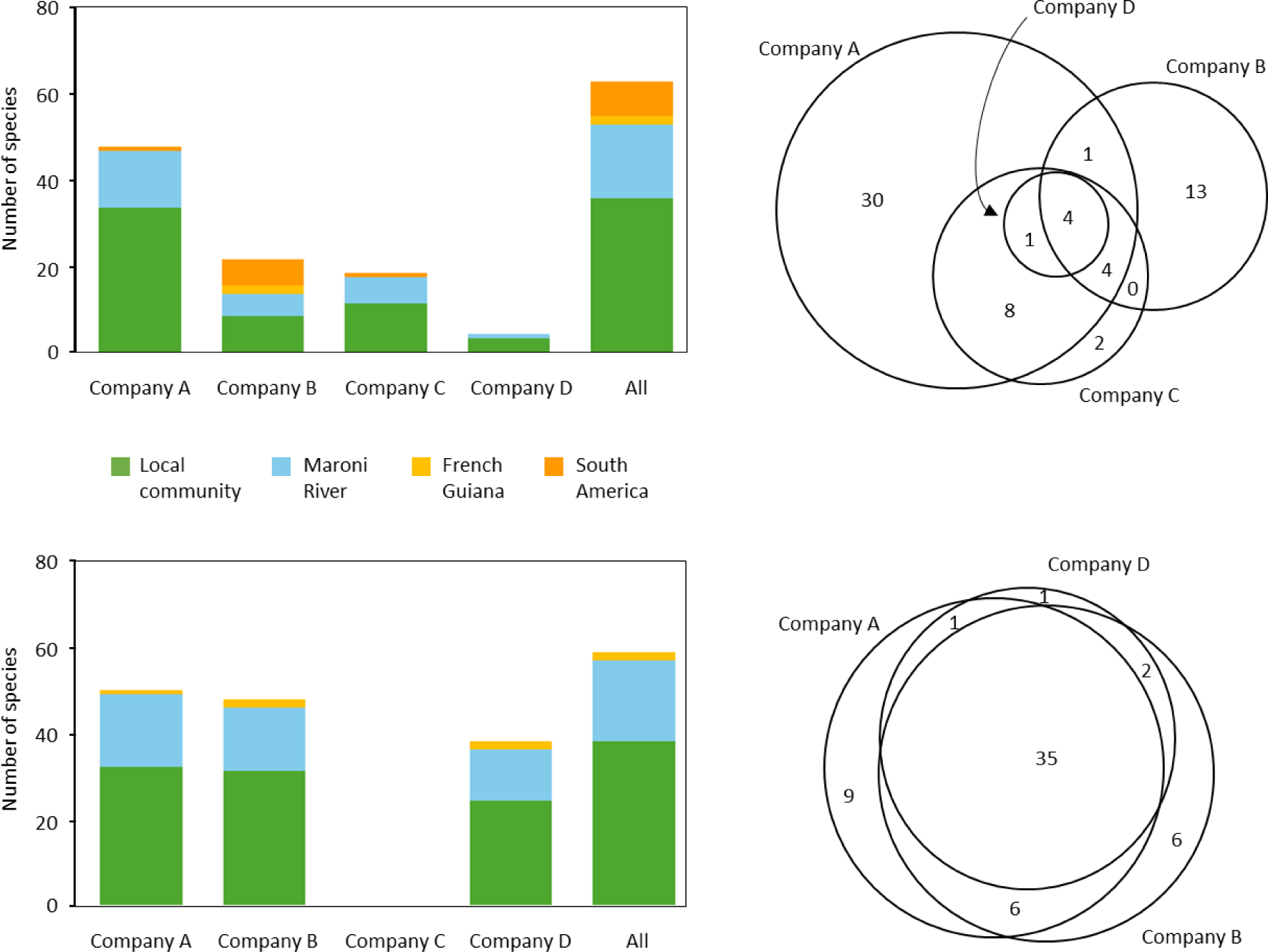
Number of fish species detected in water samples using eDNA metabarcoding based on lists provided by eDNA companies (top) and after reanalysing their raw sequencing data (bottom). Histograms show the geographic origin of the species detected: local community (*i.e.* captured on site immediately after sampling eDNA), in the Maroni River catchment, other large rivers in French Guiana, or elsewhere in South America. Venn diagrams show the number of species detections shared among companies.

Reanalysing the raw sequencing data greatly increased the number of species detected (43-52 per company) and the percentage of species in the local fish community detected (60%-67%) (Figure 1). It also decreased the differences among companies and increased the number of species detected by all companies to 34 (Figure 1). Similarly, detections of species not present in the Maroni River catchment or other large rivers in French Guiana decreased from 10 to 2 (Figure 1). See Supplementary Material S6 for the complete list of species detected after reanalysing the raw sequencing data of each company.

Among the nine species of the local fish community that were not reported by any company or detected after reanalysis of their sequencing files, only one species (*Brachyhypopomus brevirostris*) was absent from the Brosse et al. (2026) reference database. Another species, *Eleotris pisonis*, was reported by one company but was not recovered in the reanalysis, due to its absence from the Brosse et al. (2026) database.

### False positive and false negative eDNA detection rates

Comparing eDNA detections to the local fish community revealed that the companies varied greatly in their ability to detect species (Supplementary Material S7). False negative detection rates ranged from 32%-92% among the companies, whereas false positive detection rates ranged from 20.0%-59.1% (i.e. 1-14 species) (Figure 2). False negative detection rates were slightly lower for the molecular community. Differences in detection rates among the companies decreased when raw sequencing data were reanalysed, with false negative detection rates ranging from 30.0%-48.0% and false positive detection rates ranging from 32.7%-39.5% (Figure 2).

**Figure 2.**
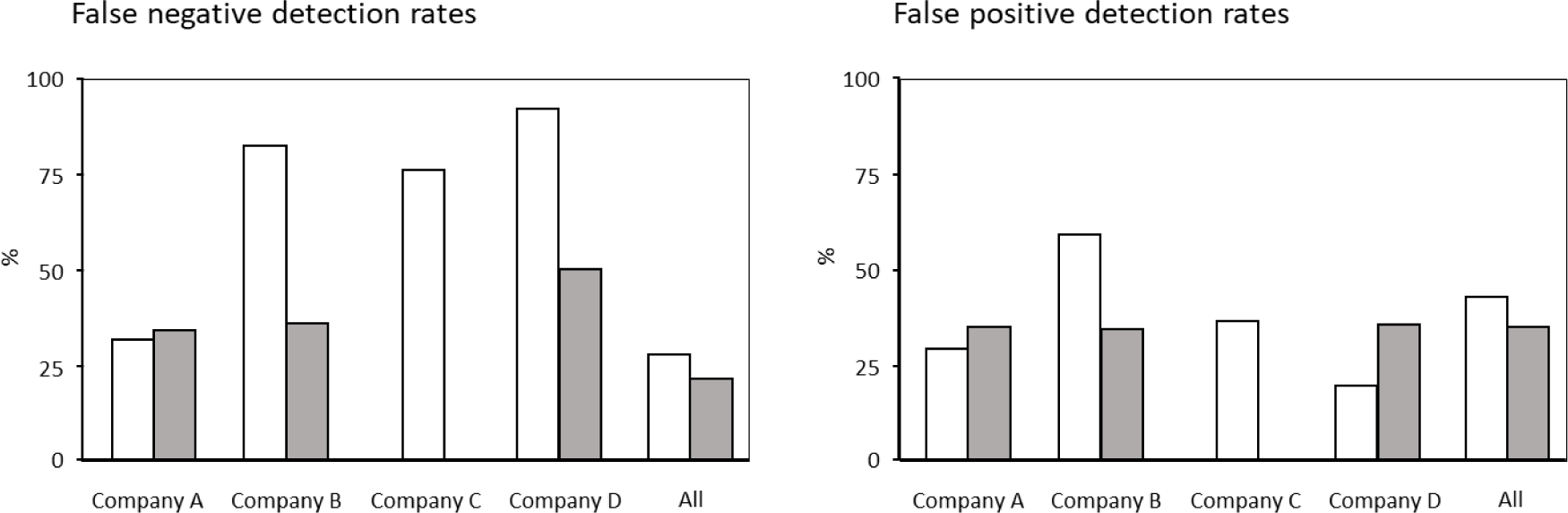
False negative detection rate (i.e. percentage of species captured in the local community not detected by eDNA barcoding) (left) and false positive detection rate (i.e. percentage of species detected by eDNA barcoding not captured in the local fish community) (right) for the inventories provided by each eDNA company and all companies combined before (white bars) and after (grey bars) reanalysing their raw sequencing data.

Reanalysing the raw sequencing data revealed that species detected by eDNA metabarcoding were significantly more abundant in the local fish community than undetected species for companies B and D (Welch’s t-test, p = 0.011 and 0.012, respectively) but not for company A (p = 0.489) (Supplementary Material S8). Similarly, detected species had significantly higher biomass than undetected species, regardless of the company (Welch’s t-test, p < 0.005) (Supplementary Material S8). We observed no significant relations between read counts of species and species abundance in the local fish community (F-test, p > 0.065), but read counts were significantly and positively correlated with species biomass (p < 0.0001) for all companies, with R² ranging from 0.31-0.59 (Supplementary Material S9).

### Fish community diversity and composition

Principal coordinate analysis showed weak dissimilarity between the fish community detected by company A and the local fish community calculated using the Jaccard index (Figure 3). The Raup-Crick index, however, revealed strong dissimilarities between the local fish community and the fish community detected by companies B, C, and D (first axis) and companies A and C (second axis) (Figure 3). The molecular community was also dissimilar from the local fish community, especially when using the Raup-Crick index.

**Figure 3.**
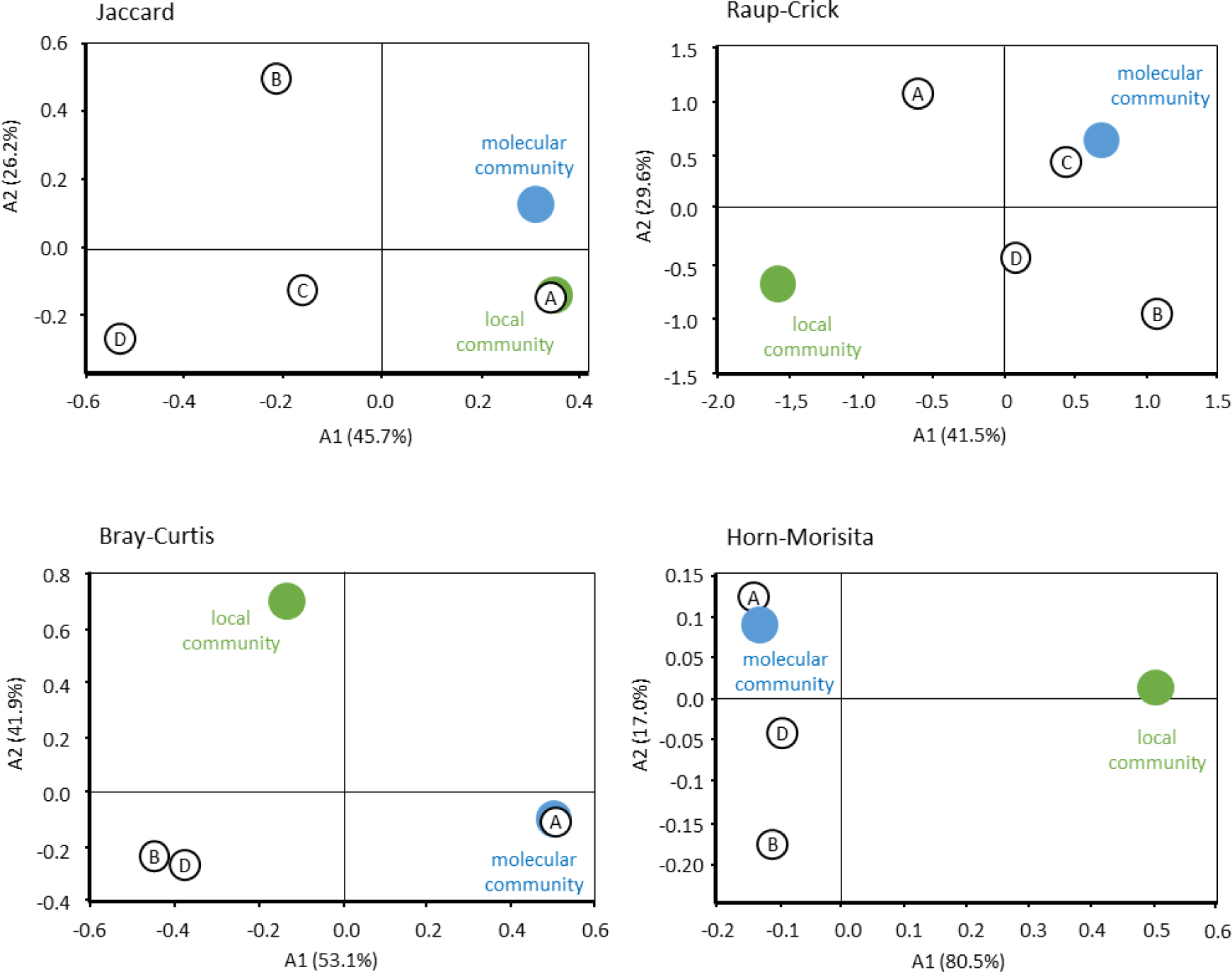
Community indices for fish captured in the local community (green circles), eDNA metabarcoding of water samples for each eDNA company (A, B, C, and D), and all companies combined (molecular community) (blue circles) on the first two axes of principal coordinate analysis (PCoA). Indices based on the presence/absence of species (Jaccard and Raup-Crick) were calculated using the commercial eDNA inventories (top). Indices based on abundance per species (Bray-Curtis and Horn-Morisita) were calculated using read counts as a proxy for abundance after reanalysing raw sequencing data (bottom).

When using read counts obtained after reanalysing the raw sequencing data as a proxy for fish abundance, we observed strong dissimilarities between eDNA metabarcoding and the local fish community along the first axis (Horn-Morisita) and second axis (Bray-Curtis) of the principal coordinate analysis, regardless of the company (Figure 3). As Company A provided the most read counts, it identified a community similar to the molecular community (Bray-Curtis and Horn-Morisita) (Figure 3).

## Discussion

### Differences among companies require caution when using eDNA metabarcoding

The companies used different protocols, from field sampling to bioinformatic analysis of DNA sequences. Without a standard protocol, non-specialist end-users may have no choice but to inventory biodiversity using eDNA metabarcoding, and the company selected could ultimately influence the quality and interpretation of the results. The eDNA inventories differed greatly in taxonomic resolution, species richness, and community composition. Although the fish species in French Guiana are well documented and have been the subject of several eDNA studies, we were surprised that many of the taxa reported were identified no finer than to genus, family, order, or even class. Only company A had satisfactory taxonomic resolution, with nearly all taxa identified to species, whereas the other three companies identified less than half of the taxa to species. Such inaccurate inventories cannot be used for conservation biology or biomonitoring, and they cast serious doubt on the ability of certain companies to provide taxonomically usable results.

In addition, the number of species in the eDNA inventories varied by a factor of 10 among companies, which is huge variability for biodiversity inventories. Reanalysing the raw sequencing data using the recently published database (Brosse et al. 2026) decreased differences among companies greatly, particularly for company D, for which the number of detected species increased from 5 to 43. This suggests that differences among companies were due mainly to differences in bioinformatic analyses rather than to those in sampling protocols or laboratory processing. Accessing a database of taxonomically certified barcodes helped decrease differences among the companies. Several eDNA studies have highlighted the importance of comprehensive and curated reference databases for detecting species reliably, as they directly influence detection accuracy, false positive and false negative detection rates, and the applicability of eDNA metabarcoding for monitoring biodiversity (Blackman et al., 2024; Weigand et al., 2019), especially for species-rich communities (Jackman et al., 2021; Keck et al., 2023; Marques et al., 2021).

Environmental managers who want to use eDNA inventories should request a guaranteed minimum degree of quality of the service that will be provided. Before subcontracting with a company to set up programmes to monitor biodiversity and environmental quality, company must be completely transparent about its methods, especially how it processes samples in the laboratory (e.g. DNA extraction method, number of PCR replicates, sequencing depth) and analyses the raw sequencing data (e.g. noise filtering, detection threshold, details of the barcode database used). Being able to access and use comprehensive databases is one of the main conditions for obtaining reliable results while respecting the FAIR principles (findability, accessibility, interoperability, and reusability). Companies should provide this information to customers, along with reviews of eDNA methods and best practices (*e.g.* Bruce et al., 2021; Dickie et al., 2018; Guillera-Arroita et al., 2017; Lahoz-Montfort et al., 2016; Pawlowski et al., 2020). Furthermore, it seems necessary to select a company that will provide the raw sequencing data, along with information about how these data are distributed in biological and technical replicates. Doing so would help to guarantee the quality of the service, as the raw sequencing data can be used to assess the accuracy of the inventory. It would also enable future assessment and updates based on changes in taxonomic knowledge, development of statistical models, and available barcode databases.

Finally, it is necessary to remain cautious when interpreting the results of eDNA inventories. For example, the companies listed 10 species that have never been observed in the Maroni River catchment (2 species) or other large rivers in French Guiana (8 species) (Supplementary Material S6). A commercial eDNA inventory should not be used to inventory biodiversity without the supervision of specialists who have taxonomic expertise in the study area.

### False negative detections highlight technical limitations of eDNA inventories

This study highlights critical issues related to interpreting eDNA inventories, which illustrates some of the limitations of this method, as in previous studies (*e.g.* Lacoursière-Roussel & Deiner, 2021; Roussel et al., 2015). Despite extensive research on eDNA in aquatic environments over the past 10 years, it is surprising that few studies have compared eDNA inventories of fish to the actual communities in the field. For example, studies that drained and restocked freshwater ponds with known fish biomasses found a moderate correlation between fish biomass and the number of reads detected in water samples per species (Di Muri et al., 2020; Ge et al., 2025). However, this type of approach is not feasible in running water, for which most studies compared eDNA metabarcoding to traditional sampling protocols (*e.g.* Gehri et al., 2021; Liu et al., 2024), which are also selective of the species, body size, and shape of fish (Cowx & Lamarque, 1990; Hamley, 1975).

The present study’s assessment of the performance of eDNA metabarcoding provides an unprecedented comparison of molecular detections to an exhaustive inventory of the local fish community in a stream. The shallow and accessible aquatic habitats in the Bastien stream allowed for such inventory (Supplementary Material S2). The electrofishing device was considered effective in previous studies of similar tropical streams (Pottier et al., 2019, 2022). Moreover, the four electrofishing passes in the study area exceeded the traditional protocol for this type of inventory (Carle & Strub, 1978; De Lury, 1951). Thus, we believe that we obtained a comprehensive inventory of the local fish community.

Among the 32%-92% of the species that the eDNA companies did not detect, three *Hemigrammus* spp. were captured via electrofishing. Although species-specific DNA markers were available for these species, only one of them was identified, and only by company B. Similarly, three species in the genera *Hyphessobrycon* and *Pristella* were captured via electrofishing but were not detected by eDNA metabarcoding. That 30%-48% of species captured via electrofishing remained undetected after reanalysing the raw sequencing data indicates that species were missing due to limitations of eDNA detection, likely caused by low DNA concentrations in the water and incomplete sampling of the environment. Indeed, among the species captured via electrofishing, those detected using eDNA metabarcoding had significantly higher biomasses than those undetected. eDNA is often considered highly effective for detecting rare and elusive species, but this result tended to demonstrate that eDNA was relatively ineffective at detecting fish species with low biomass, despite the extensive filtration in the protocols of four companies.

This raises concerns about the effectiveness of using eDNA metabarcoding alone to inventory fish biodiversity at the local scale, especially in species-rich tropical stream reaches, with rare species represented by only one or a few small individuals in traditional inventories (Pottier et al., 2019). Similar high false negative rates (up to 54%) were recently reported for eDNA metabarcoding compared to traditional netting methods in Brazilian rivers (dos Anjos Santa Rosa et al., 2025), and it is unknown whether they would be as high in temperate systems, in which eDNA is expected to persist for longer (Lamb et al., 2022). Similar studies in contrasting ecological contexts are required to better understand and quantify the extent to which eDNA metabarcoding provides false negatives, as well as potential implications for inventories of river-fish biodiversity. These studies would also benefit from using appropriate sampling designs that involve either an unambiguous inventory or relevant calibration protocols (Guillera-Arroita et al., 2017; Lahoz-Montfort et al., 2016).

### eDNA detections must be interpreted carefully

The relatively high rates of false positive detections (33%-40% among the companies) require cautious interpretation. We inventoried the local fish community by electrofishing ca. 225 m^2^ of shallow stream habitats. Given the effectiveness of this type of inventory in tropical streams (Pottier *et al.,* 2019, 2022), we believe that few species were missed. Among the false positive eDNA detections (Supplementary Table S6), previous inventories showed that tributaries of the Maroni River do contain certain species (*Acestrorhynchus falcatus, Cleithracara* gr. *maronii, Hoplerythrinus vittatus, Electrophorus* gr. *electricus, Ituglanis nebulosus, Saxatilia saxatilis*) whose presence in the study stream cannot be ruled out. DNA fragments may have drifted upstream of the electrofishing area (*e.g.* Deiner et al., 2016; Littlefair et al., 2023), been transported by predators, or come from contamination by human activities (*e.g.* Guilfoyle & Schultz, 2017; Inoue *et al*., 2023). Species that usually live in large rivers (e.g. *Hypomasticus nijsseni* and *Ageneiosus inermis*) may occasionally enter tributaries, especially during the wet season to feed or spawn, which could explain their eDNA detection.

However, some species detected using eDNA metabarcoding were highly improbable. For example, *Ageneiosus ucayalensis* and *Pygocentrus nattereri*, reported by company B, have never been reported in the Maroni River despite decades of inventories (Le Bail et al., 2026). These false positive detections likely resulted from errors in bioinformatic analysis or taxonomic assignment by the companies, as they were not replicated when we reanalysed the raw sequencing data. Indeed, if the reference database is not comprehensive, the assignment algorithm may select a phylogenetically similar species when using a minimum similarity threshold of 97% (Jackman et al., 2021). Another species, *Astyanax validus,* is geographically restricted to the upper reaches of rivers in French Guiana, including the Maroni River (Planquette et al., 1996), but its DNA was observed in the raw sequencing data of company A. Reanalysing the companies’ raw sequencing data also detected two species that had never been reported before in the Maroni River catchment (*Gymnotus* aff*. coropinae* and *Eigenmannia* cf. *pavulagem*) (Supplementary Table S6). Additional field sampling and verification of the reference sequences are needed to ensure that these detections were not caused by DNA contamination, assignment or identification errors. If they can be confirmed, they could be used to update the distribution of these species in French Guiana. These examples highlight the need for expertise on fish-species distribution at the regional and continental scales to critically interpret commercial eDNA inventories and avoid validating those that are biased. It also shows the difficulty in assessing the reliability of detection without transparent reporting of sequences and reference specimens.

These results support that detection at a given location does not automatically indicate that the species occupies the sampled habitat. This spatial ambiguity is especially problematic for large-scale biomonitoring programmes, such as those for the European Union’s Water Framework Directive, in which ecological status is expected to be based on local diversity to address the damage to biodiversity and the functioning of entire ecosystems. The standard fish-based biotic indices are based on well-defined habitats and quantitative sampling, which can be used to assess ecological degradation using robust statistical indices (Pinna et al., 2023). However, high false detection rates, either positive or negative, may strongly decrease the reliability of eDNA metabarcoding for ecological risk assessment at the local scale (Yoccoz et al., 2001). Indeed, differences between eDNA inventories and an actual community could influence the values of biotic indices. The present study demonstrates this concern. The dissimilarity indices based on presence/absence data (here, Jaccard and Raup-Crick) revealed discrepancies between eDNA-derived communities and the local community, which indicates incomplete species coverage. The performances of abundance-based indices (Bray-Curtis and Horn-Morisita) were not better when eDNA read counts were used as a proxy for species abundance. The lack of consistency between the commercial eDNA inventories and the actual fish community raises questions about using eDNA-based fish-community indices for bioindication (*e.g.* Pawlowski et al., 2018; Pont et al., 2021), at least in species-rich contexts, such as in tropical environments. As a precaution, future studies should focus on understanding the performances of eDNA-based community indices before considering using them in large-scale biomonitoring programmes and ecological risk assessments.

### Conclusion and practical recommendations

Although eDNA metabarcoding has potential for inventorying fish communities, this study indicates that its current application by eDNA companies is limited by high variability in detection rates and taxonomic accuracy. The high rates of false negatives and positives and the large differences among companies’ results highlight current limitations of eDNA metabarcoding in capturing local biodiversity in a species-rich context. This is particularly true in running water, in which animal movements and passive DNA transport further complicate spatial interpretation of the results.

Advancing eDNA metabarcoding for operational monitoring requires developing standard protocols that include field sampling, laboratory workflows, and bioinformatic analysis (Theroux et al., 2025). Using relevant sampling designs (e.g. biological and technical replicates, calibration protocols) would decrease uncertainty in the presence or absence of a taxon, and sharing databases would decrease uncertainty in taxonomic assignment. The former requires balancing the number of technical and biological replicates, as well as using blank samples for calibration to estimate false detection rates accurately. In tropical streams, future research on species-specific eDNA shedding rates and the environmental factors that influence DNA persistence and transport could help refine eDNA methods. At present, eDNA users should prioritise companies that are transparent about their methods and provide raw sequencing data based on biological and technical replicates for independent verification. Although reanalysing the raw sequencing data with a single reference database substantially reduced differences among eDNA companies, this alone cannot resolve all sources of error. Reliable eDNA inventories ultimately depend on reference databases that are curated and taxonomically certified by specialist taxonomists.

Despite the many studies that used eDNA metabarcoding and those that promoted using it for biomonitoring, it is surprising that few of them compared eDNA inventories to existing communities. This study does it for a fish community in a small, species-rich tropical stream, and we recommend increasing the number of case studies in other ecological contexts. This is a direct and rapid way to assess the performance of eDNA metabarcoding for future fish inventories and biomonitoring in running water. Based on our findings, caution is currently warranted when using only eDNA metabarcoding for regulatory assessments, especially in species-rich environments in which undetected species could bias ecological interpretations. Finally, managers who use commercial eDNA inventories to assess fish diversity should consider that interpreting them well requires expert knowledge of the taxonomy and ecology of the species in the environment studied.

## Authors’ contributions

JMR, EQ, RV and HL conceived the ideas and designed methodology; GP, RV, HL, GQ collected the data at field; PYLB, RC, GQ identified the species and standardized the names of taxa that the companies provided; BB, GL, JMR, OD and EP analysed and interpreted the data; JMR and HL led the writing of the manuscript. All authors contributed critically to the drafts and gave final approval for publication.

## Acknowledgements

We are indebted to the Direction Generale des Territoires et de la Mer (DGTM Guyane) for financial support and to Nicolas Heitz for insightful discussions on the objectives and implementation of this project.

**The bioinformatics data and pipelines used in this article are available on:**

**https://doi.org/10.5281/zenodo.19880311**

**No conflict of interest to declare.**

