## Supplementary material for "Variable performances of commercial eDNA inventories challenge their use for surveying stream fish communities": Roussel_et_al_SuppMat.pdf

**Supplementary Material S1.** Map of the Maroni River, with the Bastien stream and the location of the study site (5.27048° N, 54.234217° W).

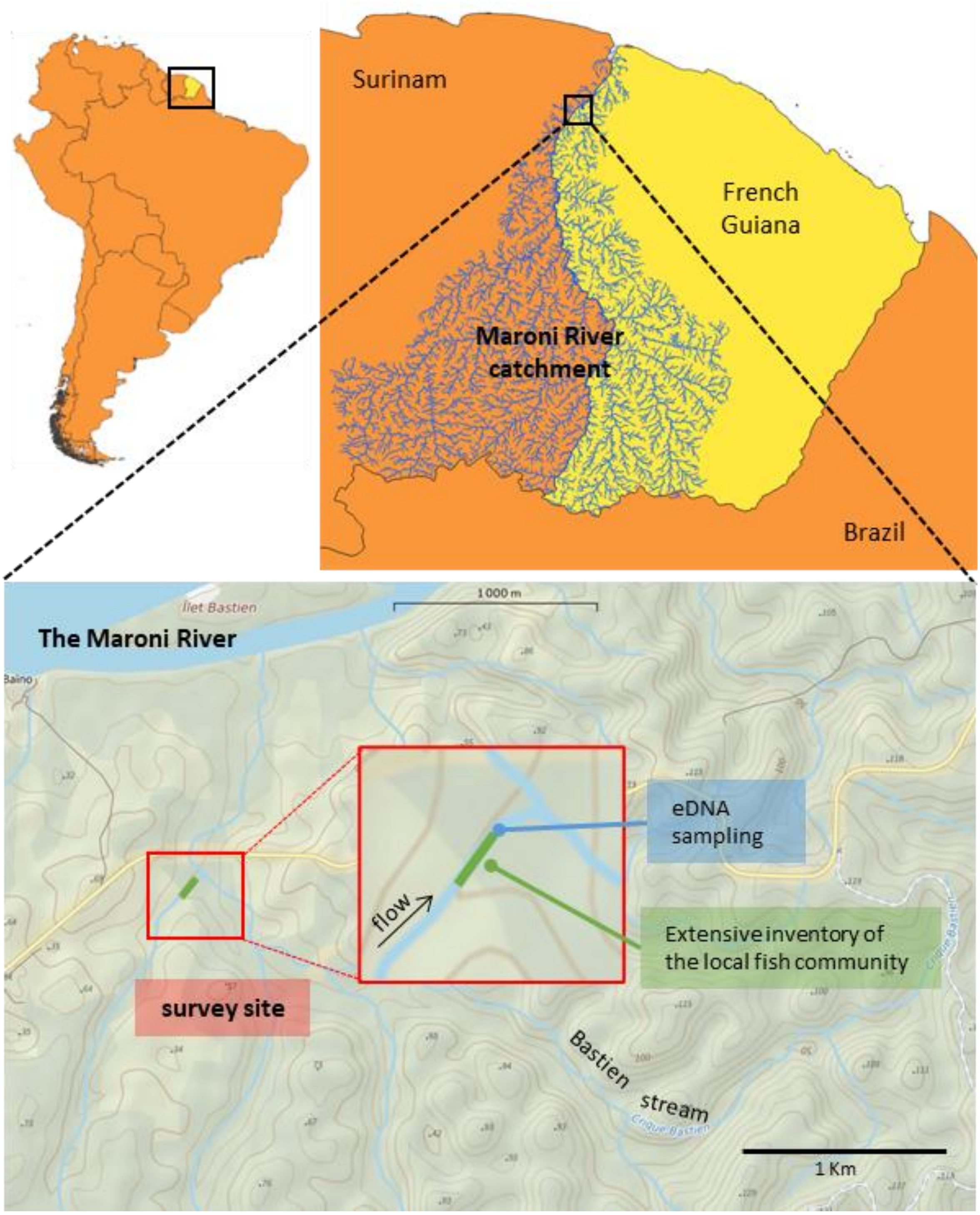

**Supplementary Material S2.** Photographs of the study site where the local fish community was inventoried. Barrier nets were installed across the channel to prevent fish from leaving or entering during the electrofishing (photographs 2 and 4).

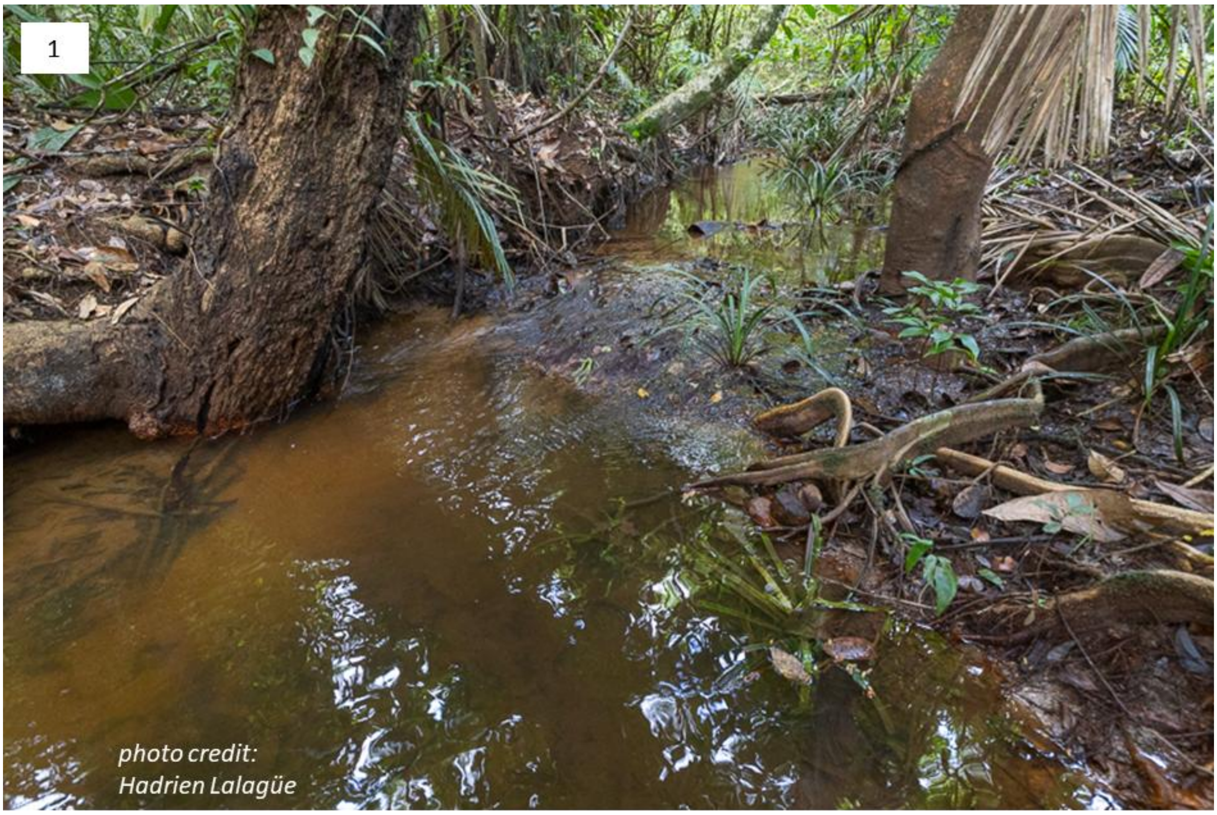


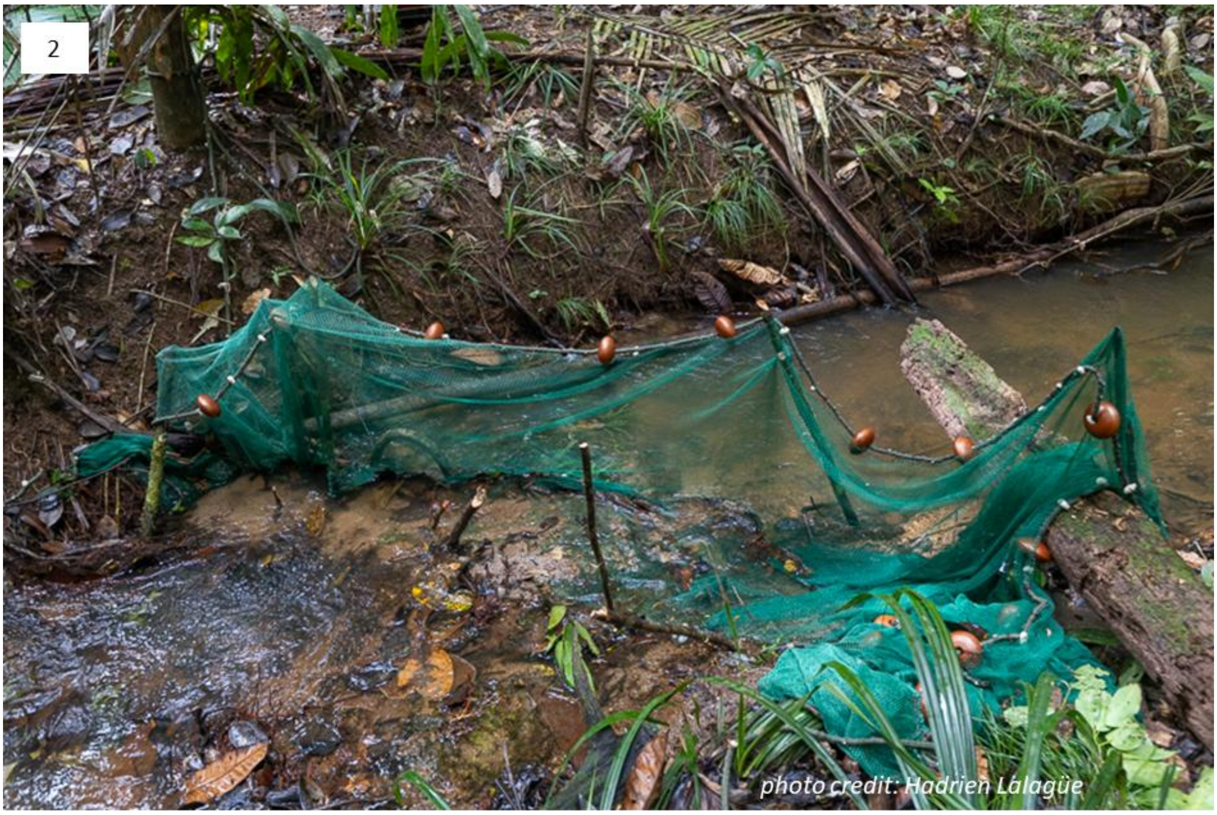

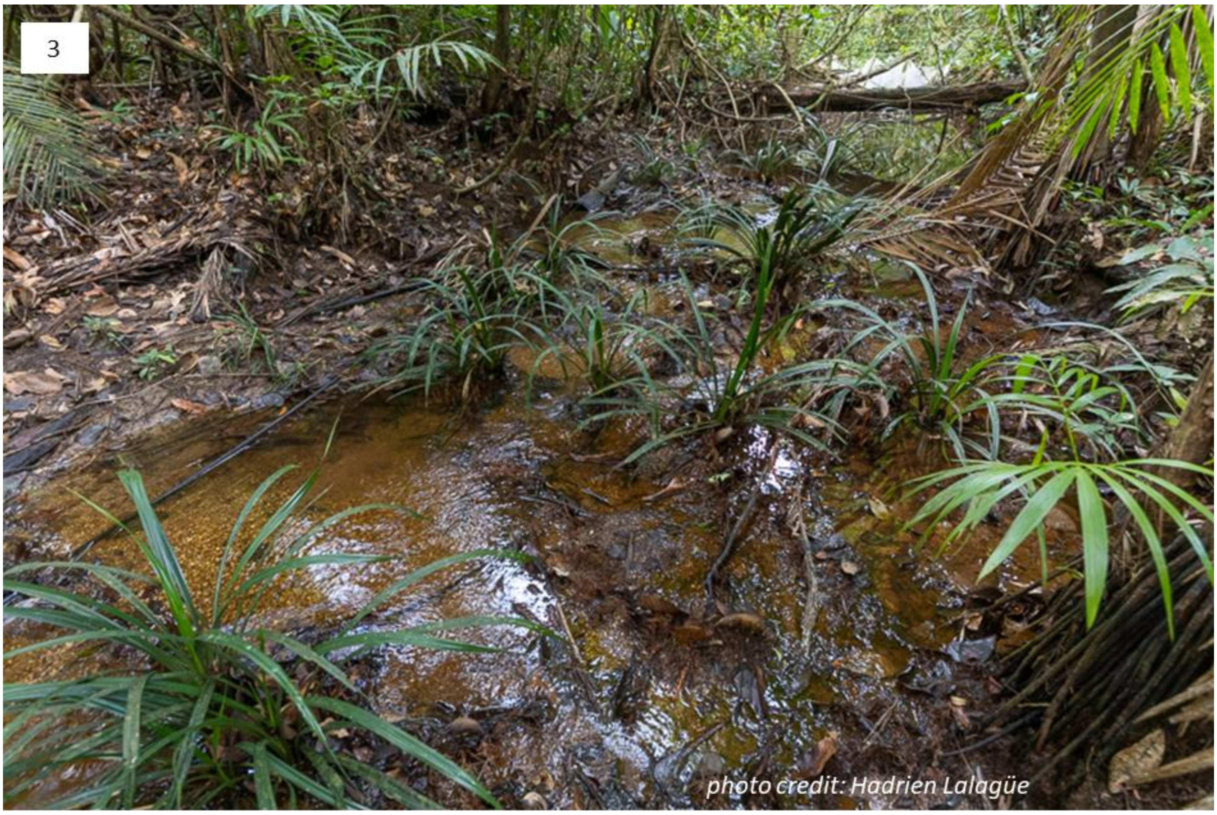
3
photo credit: Hadrien Lalagüe

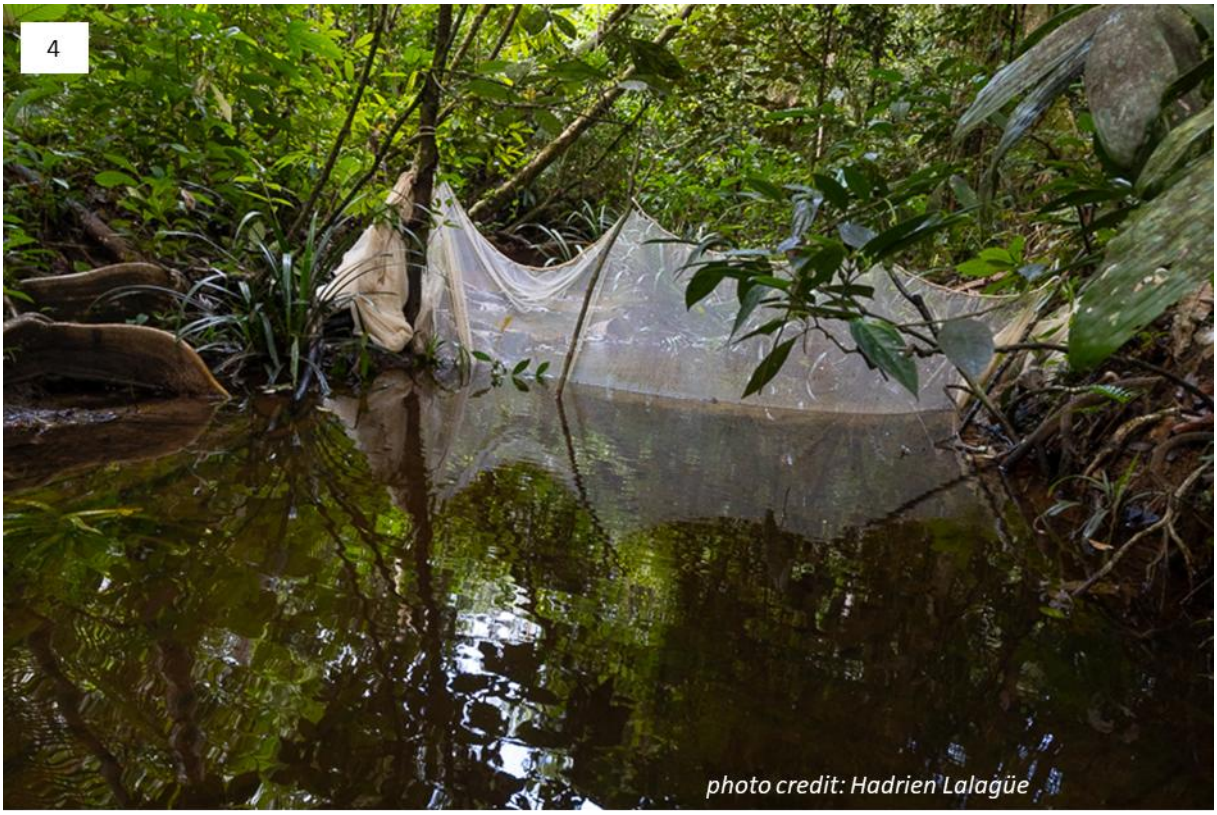
4
photo credit: Hadrien Lalagüe

**Supplementary Material S3.** eDNA sampling protocols used in the field, in the laboratory, and during bioinformatic analysis based on information provided by each eDNA company. The table also shows the number of reads and Amplicon Sequence Variants (ASVs) identified in raw sequencing data (FastQ files) when provided by the companies. PCR: polymerase chain reaction; np: information not provided by the company.

| | Company | | | |
|---|---|---|---|---|
| Characteristic | A | B | C | D |
| Number of filters (biological replicates) | 2 | 2 | 2 | 3 |
| Total volume filtered (L) | 41.5 | 39.5 | 38.5 | 3.9 |
| Total filtration time (min) | 80 | 80 | 80 | 33 |
| Filter pore size (µm) | 0.45 | 0.45 | 0.45 | 0.80 |
| PCR replicates (technical replicates per filter) | 12 | 8 | np | 1 |
| Price per sample (i.e. for 1 filter, in €) | 559 | 305 | 451 | 659 |
| Primer pair used | 12S-Teleo | 12S-MiFish | np | 12S-MiFish |
| FastQ files | Yes | Yes | No | Yes |
| Number of reads in FastQ file | 1,631,699 | 80,882 | np | 140,884 |
| Number of reads after filtering | 971,433 | 59,639 | np | 80,391 |
| Number of ASVs after filtering | 5,794 | 484 | np | 307 |
| Number of assigned reads | 925,782 | 41,160 | np | 70,077 |
| Number of assigned ASVs | 1,649 | 245 | np | 215 |

**Supplementary Material S4.** Abundance, body length (mean, standard deviation (SD), minimum, and maximum, in mm), and estimated biomass (g) per species in the local community (Bastien stream, French Guiana). The inventory resulted from electrofishing on 18 March 2024. "gr." indicates a species complex in the genus.

| Order | Family | Species or species complex (gr.) | Abundance | Body length (mm) | | Biomass (g) |
|---|---|---|---|---|---|---|
| | | | | mean (SD) | min - max | |
| Perciformes | Cichlidae | *Aequidens* gr. *tetramerus* | 2 | 97.0 (15.6) | 86 - 108 | 83.9 |
| Cyprinodontiformes | Rivulidae | *Anablepsoides* gr. *lungi* | 3 | 45.0 (4.6) | 40 - 49 | 3.8 |
| Siluriformes | Loricariidae | *Ancistrus temminckii* | 14 | 37.9 (22.9) | 17 - 79 | 43.3 |
| Characiformes | Characidae | *Bario* gr. *oligolepis* | 2 | 62.0 (22.6) | 46 - 78 | 18.1 |
| Siluriformes | Pseudopimelodidae | *Batrochoglanis raninus* | 3 | 55.0 (11.8) | 42 - 65 | 14.2 |
| Gymnotiformes | Hypopomidae | *Brachyhypopomus brevirostris* | 4 | 76.8 (54.1) | 0 - 123 | 9.6 |
| Characiformes | Characidae | *Bryconamericus* aff. *hyphesson* | 1 | 11.0 | 11 - 11 | 0.1 |
| Characiformes | Characidae | *Bryconamericus guyanensis* | 27 | 31.6 (3.3) | 25 - 36 | 14.2 |
| Characiformes | Iguanodectidae | *Bryconops affinis* | 1 | 60.0 | 60 - 60 | 3.9 |
| Siluriformes | Callichthyidae | *Callichthys callichthys* | 2 | 48.0 (0) | 48 - 48 | 5.7 |
| Characiformes | Crenuchidae | *Characidium* gr. *zebra* | 2 | 45.0 (8.5) | 39 - 51 | 2.6 |
| Siluriformes | Heptapteridae | *Chasmocranus* aff. *brevior* | 10 | 45.1 (11.5) | 20 - 60 | 6.3 |
| Siluriformes | Heptapteridae | *Chasmocranus* aff. *longior* sp1 | 1 | 40.0 | 40 - 40 | 0.5 |
| Characiformes | Lebiasinidae | *Copella* aff. *carsevennensis* sp2 | 18 | 29.6 (7.9) | 11 - 42 | 5.9 |
| Gymnotiformes | Sternopygidae | *Eigenmannia* gr. *virescens* | 2 | 98.5 (16.3) | 87 - 110 | 6.4 |
| Perciformes | Eleotridae | *Eleotris amplyopsis* | 14 | 31.7 (10.2) | 0 - 44 | 8.7 |
| Perciformes | Eleotridae | *Eleotris pisonis* | 9 | 38.3 (4.5) | 31 - 46 | 8.4 |
| Gymnotiformes | Gymnotidae | *Gymnotus* gr. *carapo* | 3 | 214.3 (85.8) | 151 - 312 | 92.8 |
| Gymnotiformes | Gymnotidae | *Gymnotus coropinae* | 10 | 93.2 (22) | 65 - 122 | 19.4 |
| Siluriformes | Cetopsidae | *Helogenes* gr. *marmoratus* | 17 | 52.9 (5.8) | 44 - 62 | 42.7 |
| Characiformes | Characidae | *Hemibrycon surinamensis* | 2 | 60.0 (0) | 60 - 60 | 9.0 |
| Characiformes | Characidae | *Hemigrammus boesemani* | 20 | 27.0 (4.3) | 16 - 36 | 6.1 |
| Characiformes | Characidae | *Hemigrammus unilineatus* | 10 | 35.4 (4.4) | 30 - 43 | 11.7 |
| Characiformes | Characidae | *Holopristis ocellifera* | 4 | 35.5 (3.8) | 30 - 38 | 4.6 |
| Characiformes | Erythrinidae | *Hoplias* gr. *malabaricus* | 8 | 134.5 (36.4) | 86 - 185 | 428.4 |
| Characiformes | Characidae | *Hyphessobrycon borealis* | 25 | 23.5 (3.1) | 15 - 30 | 5.2 |
| Characiformes | Characidae | *Hyphessobrycon simulatus* | 7 | 29.6 (3.8) | 24 - 34 | 4.1 |

| | | | | | | |
|---|---|---|---|---|---|---|
| Gymnotiformes | Hypopomidae | *Hypopomus litaniensis* | 4 | 151.3 (34.8) | 111 - 190 | 35.4 |
| Gymnotiformes | Hypopomidae | *Hypopygus* gr. *lepturus* | 5 | 67.0 (5.7) | 60 - 72 | 5.3 |
| Siluriformes | Trichomycteridae | *Ituglanis* gr. *amazonicus* | 2 | 50.5 (26.2) | 32 - 69 | 2.4 |
| Characiformes | Characidae | *Jupiaba abramoides* | 41 | 62.2 (10.9) | 40 - 84 | 263.0 |
| Perciformes | Cichlidae | *Krobia itanyi* | 14 | 53.4 (25.3) | 36 - 122 | 167.0 |
| Cyprinodontiformes | Rivulidae | *Laimosemion manaensis* | 48 | 21.5 (6.2) | 0 - 35 | 7.7 |
| Characiformes | Anostomidae | *Leporinus gossei* | 1 | 87.0 | 87 - 87 | 36.3 |
| Siluriformes | Callichthyidae | *Megalechis thoracata* | 3 | 72.0 (43.9) | 34 - 120 | 69.7 |
| Characiformes | Characidae | *Moenkhausia* gr. *collettii* | 1 | 46.0 | 46 - 46 | 2.3 |
| Characiformes | Characidae | *Moenkhausia hemigrammoides* | 3 | 33.3 (1.5) | 32 - 35 | 2.7 |
| Characiformes | Characidae | *Moenkhausia moisae* | 14 | 56.4 (11.4) | 41 - 83 | 72.7 |
| Perciformes | Cichlidae | *Nannacara anomala* | 9 | 30.9 (7.2) | 22 - 48 | 10.6 |
| Characiformes | Lebiasinidae | *Nannostomus* gr. *bifasciatus* | 10 | 32.5 (3.9) | 26 - 38 | 3.8 |
| Perciformes | Polycentridae | *Polycentrus schomburgkii* | 2 | 19.5 (2.1) | 18 - 21 | 0.6 |
| Characiformes | Characidae | *Pristella maxillaris* | 1 | 28.0 | 28 - 28 | 0.5 |
| Characiformes | Lebiasinidae | *Pyrrhulina* gr. *filamentosa* | 39 | 43.7 (18.1) | 12 - 79 | 68.4 |
| Siluriformes | Heptapteridae | *Rhamdia sebae* | 1 | 148.0 | 148 - 148 | 49.0 |
| Siluriformes | Loricariidae | *Rineloricaria* gr. *stewarti* | 1 | 70.0 | 70 - 70 | 1.2 |
| Perciformes | Cichlidae | *Saxatilia* gr. *albopunctata* | 10 | 80.0 (20.6) | 52 - 121 | 104.2 |
| Characiformes | Curimatidae | *Steindachnerina varii* | 1 | 95.0 | 95 - 95 | 21.4 |
| Gymnotiformes | Sternopygidae | *Sternopygus macrurus/sabaji* | 8 | 213.6 (62.8) | 152 - 330 | 175.6 |
| Synbranchiformes | Synbranchidae | *Synbranchus* gr. *marmoratus* | 4 | 171.3 (20.2) | 143 - 190 | 70.2 |
| Siluriformes | Auchenipteridae | *Tatia brunnea* | 3 | 50.0 (3) | 47 - 53 | 7.5 |

**Supplementary Material S5.** Histograms of individual length (top) and cumulative biomass per species (bottom) of fish in the local community captured via electrofishing.

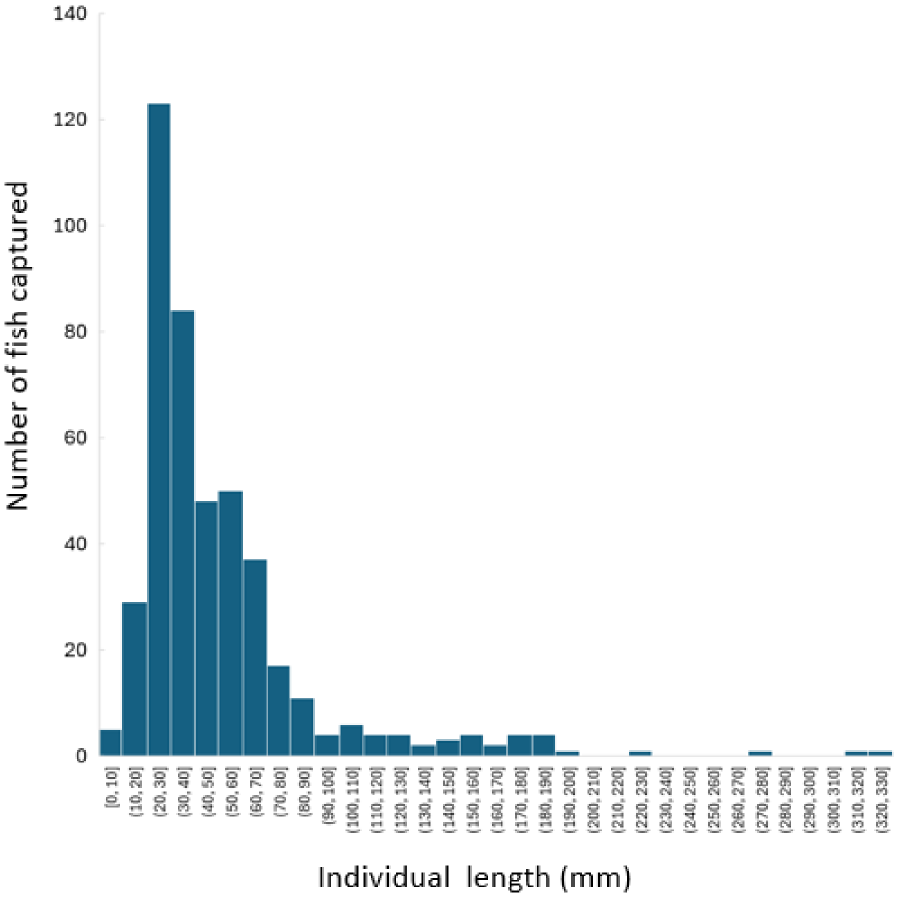


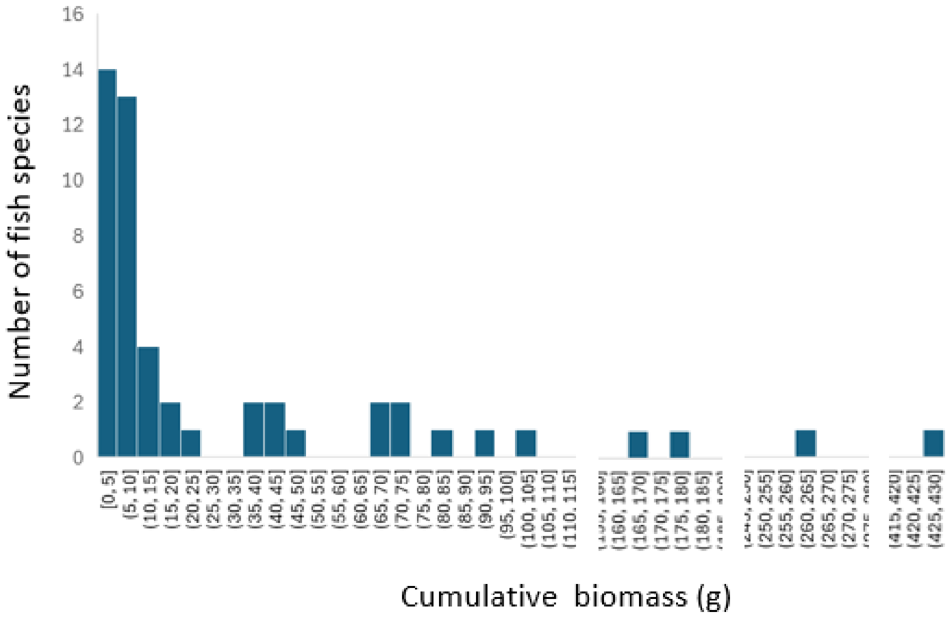

**Supplementary Material S6.** Fish species captured via electrofishing at the study site (local community) (green) in the Bastien stream (French Guiana) and detected by eDNA metabarcoding of water samples collected at the same site on the same date (18 March 2024). The table shows species reported by each eDNA company (A, B, C, and D) and those detected after this study's reanalysis of their raw sequencing data (FastQ files) (A, B, and D only). Species detected by eDNA barcoding but not captured in the local community were classified as follows: species in the Maroni River catchment (blue), in the rest of French Guiana (yellow), and in the rest of South America (brown). "gr." indicates a species complex in the genus. Barcode Index Numbers (BINs) from the Barcode of Life Data System (BOLD) are shown for each species in French Guiana. Taxonomy follows Le Bail et al. (2026), while the BINs are based on Papa et al. (2021) and Le Bail et al. (2026).

| | Species | BOLD identification | Reported by companies | Detected in FastQ files |
|---|---|---|---|---|
| South America / French Guiana / Maroni River / Local community | *Aequidens* gr. *tetramerus* | ADE2606/ADE2607 | A, C | A, B, D |
| | *Anablepsoides* gr. *lungi* | ADE2366/ADM5870 /ADM5871 | A | A |
| | *Ancistrus temminckii* | ACJ1123 | A, C | A, B |
| | *Bario* gr. *oligolepis* | ADN1547/ACV2871 | A | A, B, D |
| | *Batrochoglanis raninus* | ACE2525 | A, B, C | A, B, D |
| | *Brachyhypopomus brevirostris* | AGK0351 | | |
| | *Bryconamericus aff. hyphesson* | AEG9637 | | |
| | *Bryconamericus guyanensis* | ADH4602 | A | A, B, D |
| | *Bryconops affinis* | ADE3923 | | D |
| | *Callichthys callichthys* | AEF3201 | A, C | A |
| | *Characidium* gr. *zebra* | ADM6050/ADE2540 /ADM3248 | A | A |
| | *Chasmocranus* aff. *brevior* | ADE0651 | A | A, B, D |
| | *Chasmocranus* aff. *longior* sp1 | ADE1219 | | |
| | *Copella* aff. *carsevennensis* sp2 | ADH6438 | A | A, B, D |
| | *Eigenmannia* gr. *virescens* | ADH3981/ADH6698/ AGJ8722 | A | A |
| | *Eleotris amplyopsis* | ADK2334 | B | A, B, D |
| | *Eleotris pisonis* | AAC9395 | A | |
| | *Gymnotus* gr. *carapo* | AAB6212/AEX5853[1] | A, B, C, D | A, B, D |
| | *Gymnotus coropinae* | ADE2268 | | |
| | *Helogenes* gr. *marmoratus* | ADE2445/ADE2444/ ADE2443/AEZ0831 | | |
| | *Hemibrycon surinamensis* | ADE0487 | A | A, D |
| | *Hemigrammus boesemani* | AAB8134 | | |
| | *Hemigrammus unilineatus* | ACR8448 | B | B |
| | *Holopristis ocellifera* | ACS3953 | | A, B |
| | *Hoplias* gr. *malabaricus* | AFU2064[2]/AFV7073[3] | A, B, C, D | A, B, D |
| | *Hyphessobrycon borealis* | ADE3322 | | B, D |
| | *Hyphessobrycon simulatus* | ACR7942 | | B |
| | *Hypopomus litaniensis* | ADE1884 | A, C | A, B, D |
| | *Hypopygus* gr. *lepturus* | ADE1262/ADE1045 | A, C | A, B, D |
| | *Ituglanis* gr. *amazonicus* | ADM4536/ADE2267 | A, B | A, B |

| | | | |
|---|---|---|---|
| *Jupiaba abramoides* | ACR7858 | A | A, B, D |
| *Krobia itanyi* | ADE1084 | A | A, B, D |
| *Laimosemion manaensis* | ADE2032 | A | A, B, D |
| *Leporinus gossei* | AAC5571 | A | A, B |
| *Megalechis thoracata* | ACR8270 | A, B, C, D | A, B, D |
| *Moenkhausia* gr. *collettii* | ACR8022/ADE0784 | | |
| *Moenkhausia hemigrammoides* | ACR7727 | | |
| *Moenkhausia moisae* | ACR8418 | A | A, B, D |
| *Nannacara anomala* | AED8776 | A, B, C, D | A, B, D |
| *Nannostomus* gr. *bifasciatus* | ADH3809/ADH3810/ AEX8086 | A | A, B, D |
| *Polycentrus schomburgkii* | AAY2847 | A | B |
| *Pristella maxillaris* | AAC9546 | | |
| *Pyrrhulina* gr. *filamentosa* | ADE1661/ADE1584 | A | A, B, D |
| *Rhamdia sebae* | ADE4073 | A, B, C | A, B, D |
| *Rineloricaria* gr. *stewarti* | ACJ0428/ADH8611 | A | A, B |
| *Saxatilia* gr. *albopunctata* | ADJ6307/ADE2074/ ADH4408 | A | A, B, D |
| *Steindachnerina varii* | ACR8536 | | |
| *Sternopygus macrurus/sabaji* | ADH7013/ACG7687 | A, C | A, B, D |
| *Synbranchus* gr. *marmoratus* | ADM6000/ADE2961 | A | A, B, D |
| *Tatia brunnea* | ACR8870 | A | A |
| *Acestrorhynchus falcatus* | ABU8139[4] | A | A |
| *Ageneiosus inermis* | AAC6222[5] | | B |
| *Ancistrus* aff. *hoplogenys* | ACJ1201/ACJ1296 | | A, B |
| *Astyanax bimaculatus* | AAA7655 | B | |
| *Astyanax validus* | ACJ1803 | A | A |
| *Cetopsidium* gr. *orientale* | ADE4127/ADH4823/ ADH4822 | A | A |
| *Cleithracara* gr. *maronii* | ADM3513/ADE2633 /ADH3965 | A | A, B, D |
| *Electrophorus gr. electricus* | ACA1272/ACE2732 | A, B, C | A, B, D |
| *Erythrinus erythrinus* | ADE1155 | A, B, C | A, B |
| *Gymnotus* gr. *anguillaris* | ADH4414/ADE2269 | A | A, B, D |
| *Heptapterus bleekeri* | | | B, D |
| *Hoplerythrinus vittatus* | ADC3975 | A, C, D | A, B, D |
| *Hypomasticus nijsseni* | ACF5028 | A | A, B, D |
| *Ituglanis nebulosus* | ADM6925 | A | A, B, D |
| *Japigny kirschbaum* | ACF8576 | A, C | A, B, D |
| *Krobia guianensis* | ADE1186 | B | A |
| *Microcharacidium eleotrioides* | ADE1286 | A | A, B, D |
| *Moenkhausia* aff. *surinamensis* | ACV2805 | A, C | A, B, D |
| *Saxatilia saxatilis* | AAE4421 | C | A, B, D |
| *Trachelyopterus* aff. *maculosus* | AAC7401 | B | A, B, D |
| *Ageneiosus ucayalensis* | AAC6221 | B | |
| *Eigenmannia* cf. *pavulagem* | ACC5102 | | A, B, D |

| | | | |
|---|---|---|---|
| *Gymnotus* aff. *coropinae* | AGJ9856 | | B, D |
| *Pygocentrus nattereri* | ABZ7351 | B | |
| *Ancistrus* sp. 2a MNRJ42890 | NA | A | |
| *Eigenmannia macrops* | NA | B | |
| *Gymnotus sylvius* | NA | C | |
| *Hemigrammus pretoensis* | NA | B | |
| *Hyphessobrycon megalopterus* | NA | B | |
| *Saxatilia inpa* | NA | B | |
| *Synbranchus* sp. NS-2021 | NA | B | |
| *Trichomycterus areolatus* | NA | B | |

[1]AAE3544; [2]ABZ3047; [3]ACF3787; [4]ADH6781;[5]ADZ9108 in Papa et al. (2021)


**Supplementary Material S7.** Number of species detected by each eDNA company and/or captured during the extensive inventory of the local community via electrofishing. The false negative and positive detection rates are shown. Numbers in square brackets are the values obtained after reanalysing the companies' raw sequencing data, when available.

| | Company | | | | |
|---|---|---|---|---|---|
| **Result** | **A** | **B** | **C** | **D** | **All** |
| Detected and captured | 34<br>*[33]* | 9<br>*[32]* | 12 | 4<br>*[25]* | 36<br>*[39]* |
| Detected but not captured | 14<br>*[18]* | 13<br>*[17]* | 7 | 1<br>*[14]* | 27<br>*[21]* |
| Not detected but captured | 16<br>*[17]* | 41<br>*[18]* | 38 | 46<br>*[25]* | 14<br>*[11]* |
| False-negative detection rate | 32%<br>*[34%]* | 82%<br>*[36%]* | 76% | 92%<br>*[50%]* | 28%<br>*[22%]* |
| False-positive detection rate | 29.2%<br>*[35.3%]* | 59.1%<br>*[34.7%]* | 36.8% | 20%<br>*[35.9%]* | 42.9%<br>*[35.0%]* |

**Supplementary Material S8.** Relation between the eDNA detection of species (yes or no) and the abundance (top) and biomass (bottom) of fish species in the local community (electrofishing), for the three companies that provided raw sequencing data. The p-values are based on Welch's t-tests (unequal variance, two samples).

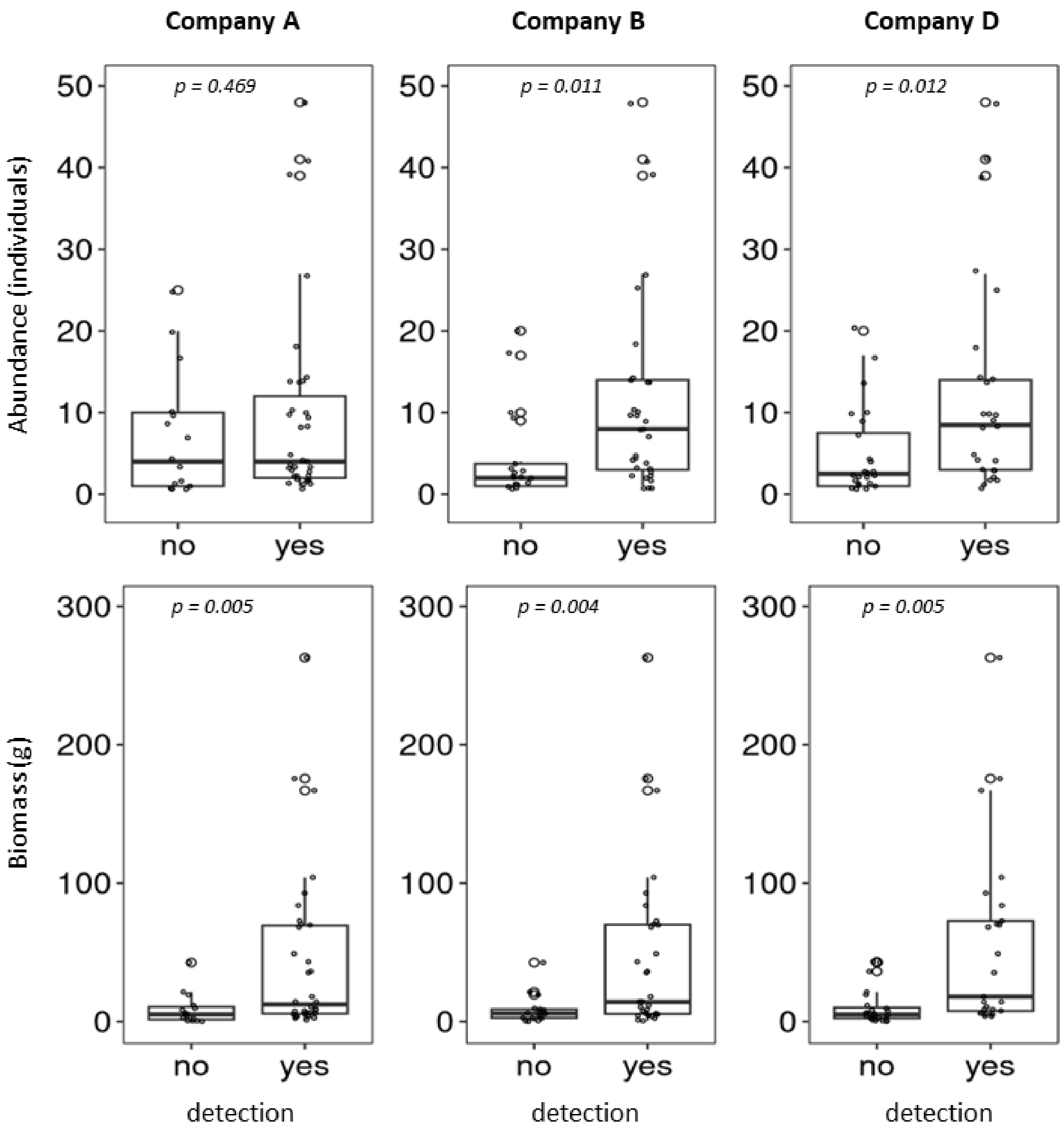

**Supplementary Material S9.** Relation between the abundance (top) and biomass (bottom) of fish species in the local community (electrofishing) and the number of reads for the corresponding species for eDNA inventories of the three companies that provided raw sequencing data. The graphs show linear regressions, 95% confidence intervals (grey shading), and adjusted R-squared and p-values based on the F-test.

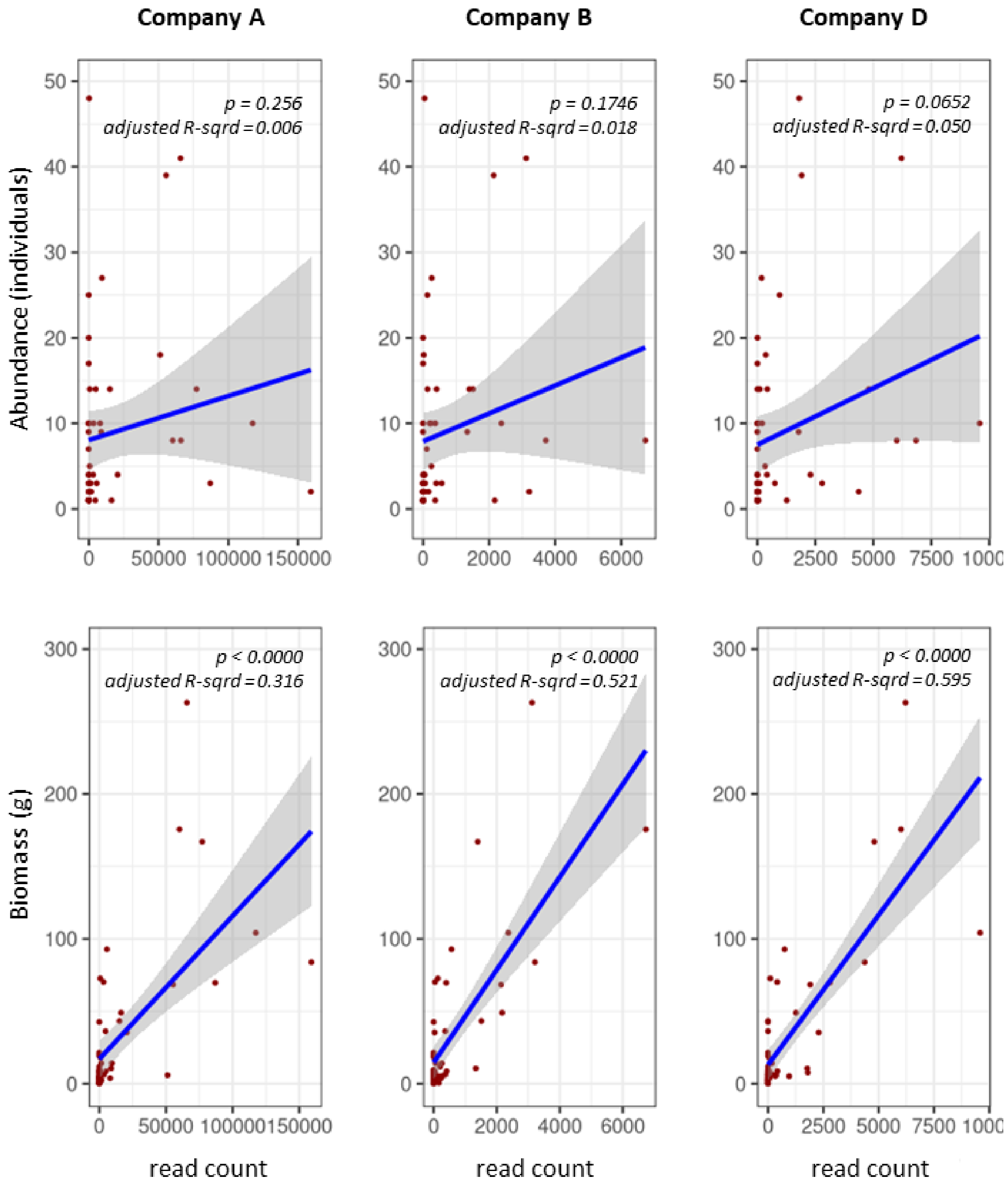
